# The toll-like receptor 7 agonist imiquimod increases ethanol self-administration and induces expression of toll-like receptor related genes

**DOI:** 10.1101/2021.10.25.465772

**Authors:** Dennis F. Lovelock, Wen Liu, Sarah E. Langston, Jiaqi Liu, Kalynn Van Voorhies, Kaitlin A. Giffin, Ryan P. Vetreno, Fulton T. Crews, Joyce Besheer

## Abstract

**Background:** There is growing evidence that immune signaling may be involved in both the causes and consequences of alcohol abuse. Toll-like receptor (TLR) expression is increased by alcohol consumption and is implicated in AUD, and specifically TLR7 may play an important role in ethanol consumption.

**Methods:** We administered the TLR7-specific agonist imiquimod in male and female Long-Evans rats to determine 1) gene expression changes in brain regions involved in alcohol reinforcement, the nucleus accumbens core and anterior insular cortex, in rats with and without an alcohol history, and 2) whether TLR7 activation could modulate operant alcohol self-administration.

**Results:** Interferon regulatory factor 7 (IRF7) was dramatically increased in both sexes at both 2 and 24 h post-injection regardless of alcohol history, while TLR3 and 7 gene expression changes were region- and sex-specific. The pro-inflammatory cytokine TNFα was increased 24h post-injection in rats with an alcohol self-administration history but this effect did not persist after four injections, suggesting molecular tolerance. In both males and females, ethanol consumption was increased 24 h after imiquimod injections with sex-specific differences: in females this effect emerged following the first injection but in males this increase did not occur until the third injection, suggesting sex differences in adaptation to repeated TLR7 activation. Notably, imiquimod reliably induced weight loss, indicating that sickness behavior persisted across repeated injections.

**Conclusion:** These findings show that TLR7 activation can modulate alcohol drinking in an operant self-administration paradigm, and suggest that TLR7 and IRF7 signaling pathways may be a viable druggable target for treatment of AUD.

## Background

There is growing evidence that immune signaling is involved in both the development and persistence of alcohol use disorder (AUD; Coleman and Crews, 2018). Increased expression of proinflammatory genes such as Toll-like receptors (TLR) and cytokines have been found in the brains of postmortem alcoholics [2-4] and chronic alcohol consumption is associated with increased levels of cytokines in circulation [5]. In animal models, alcohol has been shown to increase expression of neuroimmune signaling molecules in the brain [6-9], which have in turn been shown to increase voluntary alcohol consumption [10-13]. Thus, neuroimmune signaling may serve as a positive feedback loop in AUD, being both a cause and consequence of excessive alcohol intake [1].

Toll-like receptors (TLRs) are a major component of immune signaling that were discovered to mediate responses to pathogens and necrotic cell damage. For example, TLR4 was first discovered to respond to bacterial endotoxin, and endosomal TLR3 and 7 respond to double and single-stranded RNA, respectively, that respond to viruses. The brain is generally sterile, that has led to recent studies of endogenous TLR agonist signaling in brain contributing to induction of neuronal TLR [9]. Activation of TLRs leads to signaling through adapters, MyD88 (all TLR except TLR3) and/or TRIF (TLR3 and 4), ultimately resulting in nuclear translocation and induction of pro-inflammatory genes including cytokines and interferons [4]. TLRs also have important roles in normal brain functioning. For example, TLR3, 7 and 8 regulate axonal growth, dendritic pruning, and neuronal morphology [14, 15]. Thus, TLR signaling contributes to neurobiology as well as their known roles in the immune system.

Although studies have found increased expression of multiple TLRs in post-mortem human brain of AUD patients [16], preclinical studies indicate a complex relationship with alcohol drinking. An initial study linking TLRs to alcohol consumption found that the TLR4 agonist and bacterial endotoxin LPS increased voluntary intake of alcohol in a two-bottle choice model in multiple mouse strains [17]. Interestingly, other studies found that TLR4 knockout and antagonism had little effect on ethanol consumption across both rats and mice [18, 19], suggesting that other immune factors contribute to the increase in drinking. TLR3 knockout male mice consumed less alcohol in a two-bottle choice paradigm [20], and recent studies have shown that TLR3 activation is capable of driving alcohol consumption in male rats [13] and mice [10, 11]. TLR7 was recently shown to drive voluntary ethanol consumption as mice given 10 injections of the TLR7/8 small molecule agonist R848, which only activates TLR7 in rodents [21], subsequently drank more alcohol in a two-bottle choice procedure [12]. Voluntary alcohol drinking may assess different circuitry as compared to operant self-administration, with the latter better assessing the rewarding properties of alcohol [22]. Nonetheless, work from our lab showed that TLR3 activation produced a similar increase in operant alcohol self-administration and rapid increases in TLR3 gene expression in the insular cortex and nucleus accumbens [13], a brain circuit known to be involved in the regulation of alcohol drinking and interoceptive sensitivity to alcohol [23, 24]. Further, these increases in TLR3 gene expression were positively correlated with TLR7 gene expression in the insular cortex, but not the nucleus accumbens. Thus, TLR agonists induce increases in expression of multiple TLR receptors adding to the complexity of interpreting responses to TLR specific agonists.

While initial studies examining TLR signaling and alcohol tended to focus solely on males, when female subjects have been included sex differences have been found. TLR3 activation via poly(I:C) in female mice did not increase alcohol consumption as has been seen in males, and the time course of the subsequent immune response differs with females showing a delayed response [10, 11]. Interestingly, both sexes displayed decreased drinking in TLR2 knockout mice while only MyD88 knockout males increased voluntary alcohol consumption [19]. At present, the effects of TLR7 modulation in females has yet to be examined. TLR7 makes for a particularly interesting target in females as it is located on the X chromosome and escapes X-inactivation in immune cells and thus may be more highly expressed in females at baseline [25, 26]. As such, the goals of the present study, were to examine 1) gene expression changes in the nucleus accumbens core (AcbC) and anterior insular cortex (AI) in alcohol naïve male and female Long Evans rats following administration of the TLR7 agonist imiquimod, and 2) whether imiquimod modulates alcohol self-administration in male and female rats and the consequences of multiple imiquimod injections on self-administration and gene expression. Gene expression analyses focused on TLR signaling pathways in the AcbC and AI to highlight downstream signaling and immune molecules that may play an important role in the TLR7-mediated increase in alcohol consumption.

## METHODS

### Animals

Adult male and female Long Evans rats (Envigo-Harlan, Indianapolis, IN) were delivered at 7 weeks old and were handled daily for 1 week prior to the start of the experiment. All rats were doubled housed in ventilated cages in same-sex pairs. Rats had ad-libitum access to food and water in the home cage unless noted. The rats were kept in a temperature and humidity-controlled colony room that ran on a 12-hour light/dark cycle (lights on at 07:00). All experiments were conducted during the light cycle. Animals were under the care of the veterinary staff from the Division of Comparative Medicine at UNC-Chapel Hill. All of the procedures followed the guidelines established by the NIH Guide to Care and Use of Laboratory Animals and institutional guidelines.

### Drugs

Ethanol (95% v/v; Pharmco-AAPER, Shelbyville, KY) was diluted in distilled water. Imiquimod (Sigma-Aldrich CO. LLC, Saint Louis, MO, USA; Lot# 25236) was dissolved in 45% hydroxypropyl-beta-cyclodextrin (Acros Organics, Geel, Belgium), which was also used for control injections. Imiquimod was injected intraperitoneal (IP) at a volume of 1 ml/kg.

### Apparatus

Self-administration chambers (Med Associates Inc., St. Albans, VT) were individually located within sound attenuating chambers with an exhaust fan to circulate air and mask outside sounds. Chambers were fitted with a retractable lever on the opposite walls (left and right) of the chamber. There was a cue light above each lever and liquid receptacles in the center panels adjacent to both levers. Responses on the left (i.e., active) lever resulted in cue light illumination, stimulus tone, and delivery of 0.1 ml of solution across 1.66 seconds via a syringe pump into the left receptacle once the response requirement was met. Responses on the right (inactive) lever had no programmed consequence. The chambers also had infrared photobeams which divided the floor into 4 zones to record general locomotor activity throughout each session.

### EtOH Self-Administration Training

Self-administration sessions (30 minutes) took place 5 days per week (M-F) with the active lever on a fixed ratio 2 schedule of reinforcement such that every second response resulted in delivery of EtOH [27]. A sucrose-fading procedure was used in which EtOH was gradually added to the 10% (w/v) sucrose solution. The exact order of exposure was as follows: 2% (v/v) EtOH/10% (w/v) sucrose, 2E/ 10S, 5E/10S, 10E/10S, 10E/5S, 15E/5S, 15E/2S, 20E/2S, 20E, 15E. Following sucrose fading, unsweetened EtOH/15% (w/v) was the reinforcer for the remainder of the study. At the end of each session, wells were inspected to ensure that rats had consumed all fluid.

### Brain tissue collection and sectioning

Brains were rapidly extracted and flash frozen with isopentane (Sigma-Aldrich, MI). Brains were stored at -80°C until brain region sectioning. Brains were sectioned on a cryostat (−20°C) up to a predetermined bregma coordinate for each region of interest (ROI). Then, a micropunch tool was used to collect tissue specific to each brain region. ROIs were separated by left and right hemispheres, and all real-time RTPCR experiments used the right hemisphere when separated. Brain tissue was stored at -80°C until real-time RTPCR analysis. For all experiments where tissue was collected 24h-post imiquimod injection (experiments 2-4), spleens were weighed as an index of the inflammatory response.

### Tissue processing and real-time RTPCR

RNA was extracted from brain tissue using the RNeasy Mini Kit (Qiagen, Venlo, Netherlands) according to the manufacturer’s instructions. RLT lysis buffer containing β-mercaptoethanol (Sigma Aldrich) was used for tissue homogenization. RNA concentration and purity for each sample were determined using a spectrophotometer (Nanodrop 2000, ThermoScientific). RNA was reverse transcribed into cDNA using the either the QuantiNova Reverse Transcription Kit (Qiagen) or Superscript III First-Strand Synthesis System (Invitrogen, Waltham, MA, USA) according to the manufacturer’s instructions. Following reverse transcription, all samples were diluted 1:10 with nanopure water (200 uL total) and stored at -20°C before RT-PCR experiments. Real-time RTPCR was conducted using a QuantStudio3 (ThermoFisher) for all experiments. Using a 96-well plate, each sample was run in triplicate using 10 µL total volume per well with the following components: PowerUp Sybr green dye (ThermoFisher, containing ROX dye for passive reference), forward and reverse primers (Eton Biosciences Inc., NC, USA), and cDNA template. The PCR was run with an initial activation for 10 mins at 95°C, followed by 40 cycles of the following: denaturation (95°C for 15s), annealing (60°C for 30s), and extension (72°C for 45s). Melt curves were obtained for all experiments to verify synthesis of a single amplicon. All primer sequences are displayed in Table 1.

**Table 1:**
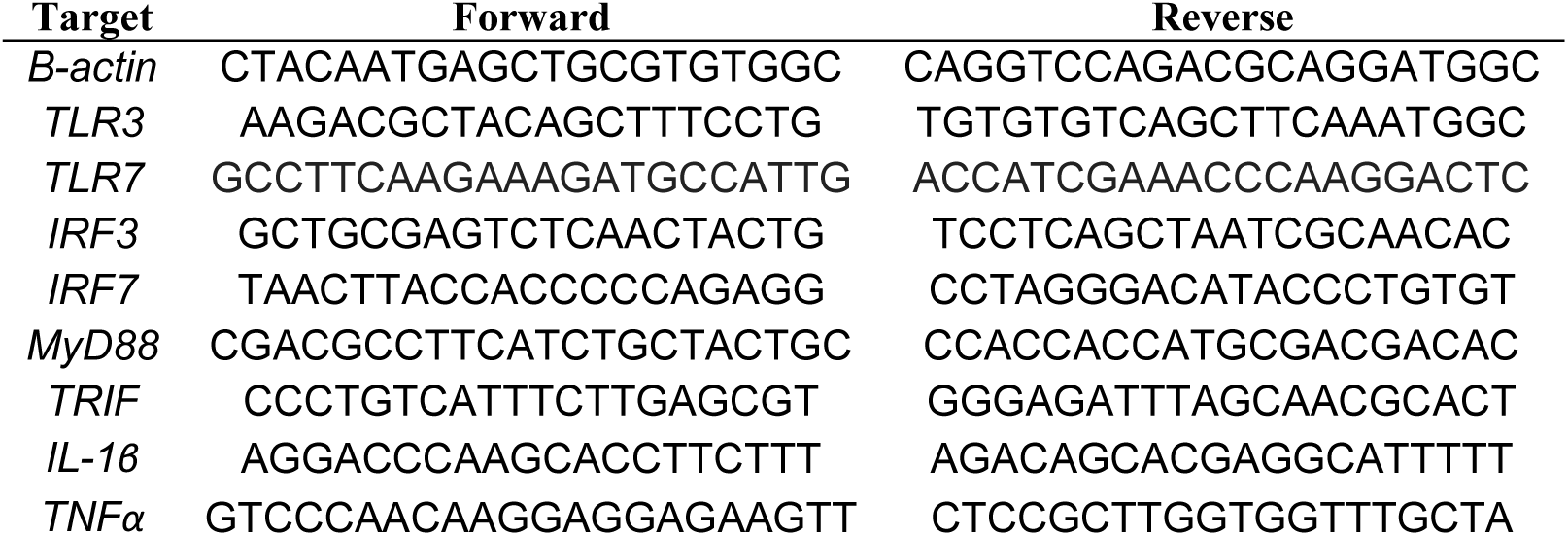
Primers used for PCR Analysis.

### Experiments

#### Experiment 1: Central gene expression 2 h post-imiquimod injection

Male and female naive rats were injected with imiquimod or vehicle (0 or 10 mg/kg, IP; n=8/dose/sex). To examine the short-term consequences of imiquimod, rats were sacrificed 2 hours post-injection (figure 1A). Brain tissue was collected for RT-PCR analysis. Spleen weights were not collected for this experiment as they were for the other experiments as they were not expected to differ 2 h post-imiquimod.

**Figure 1:**
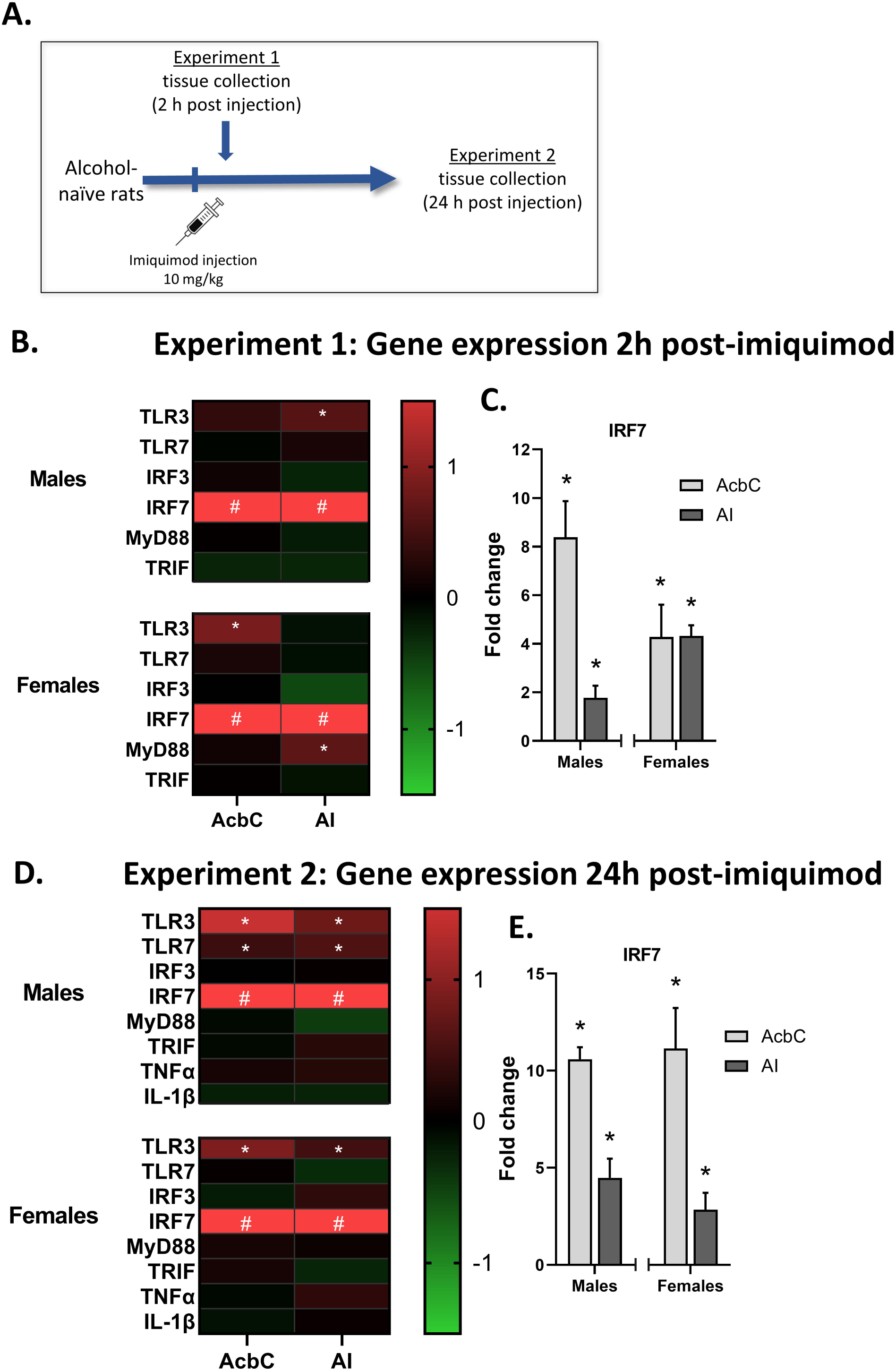
Experiment 1 and 2 gene expression. **A)** Rats were given an injection of either imiquimod (3 mg/kg) or vehicle and tissue was collected either 2h (Experiment 1) or 24h (Experiment 2) post-injection. **B)** Experiment 1 gene expression changes in the nucleus accumbens core (AcbC) and anterior insula (AI) induced by imiquimod relative to controls visualized as a heatmap. **C)** As IRF7 changes were much greater than seen in other genes these data are shown separately as fold change from vehicle controls (where vehicle group = 0 fold change). **D)** Experiment 2 gene expression changes visualized as a heatmap. At 24h post- injection TLR3 was reliably increased across both brain regions and sexes. **E)** IRF7 gene expression remained high 24h post-injection. *p<0.05 compared to vehicle control, # p<0.05 compared to control and value is greater than the heatmap scale.

#### Experiment 2: Central gene expression 24 h post-imiquimod injection

The design of this experiment was identical to that of Experiment 1 except that rats (n=8/dose/sex) were sacrificed 24 hours post-injection and spleen weights were collected as an index of the inflammatory response (figure 1A).

#### Experiment 3: Effect of EtOH self-administration history on gene expression 24 h after a single imiquimod injection

After daily self-administration of 15% ethanol for 24 days, male and female Long-Evans rats were injected with imiquimod (0 or 10 mg/kg, IP; n=10/dose/sex) and sacrificed 24 hours post-injection (figure 2A). Brains and tissue were collected for RT-PCR analysis and spleen weights were recorded.

**Figure 2:**
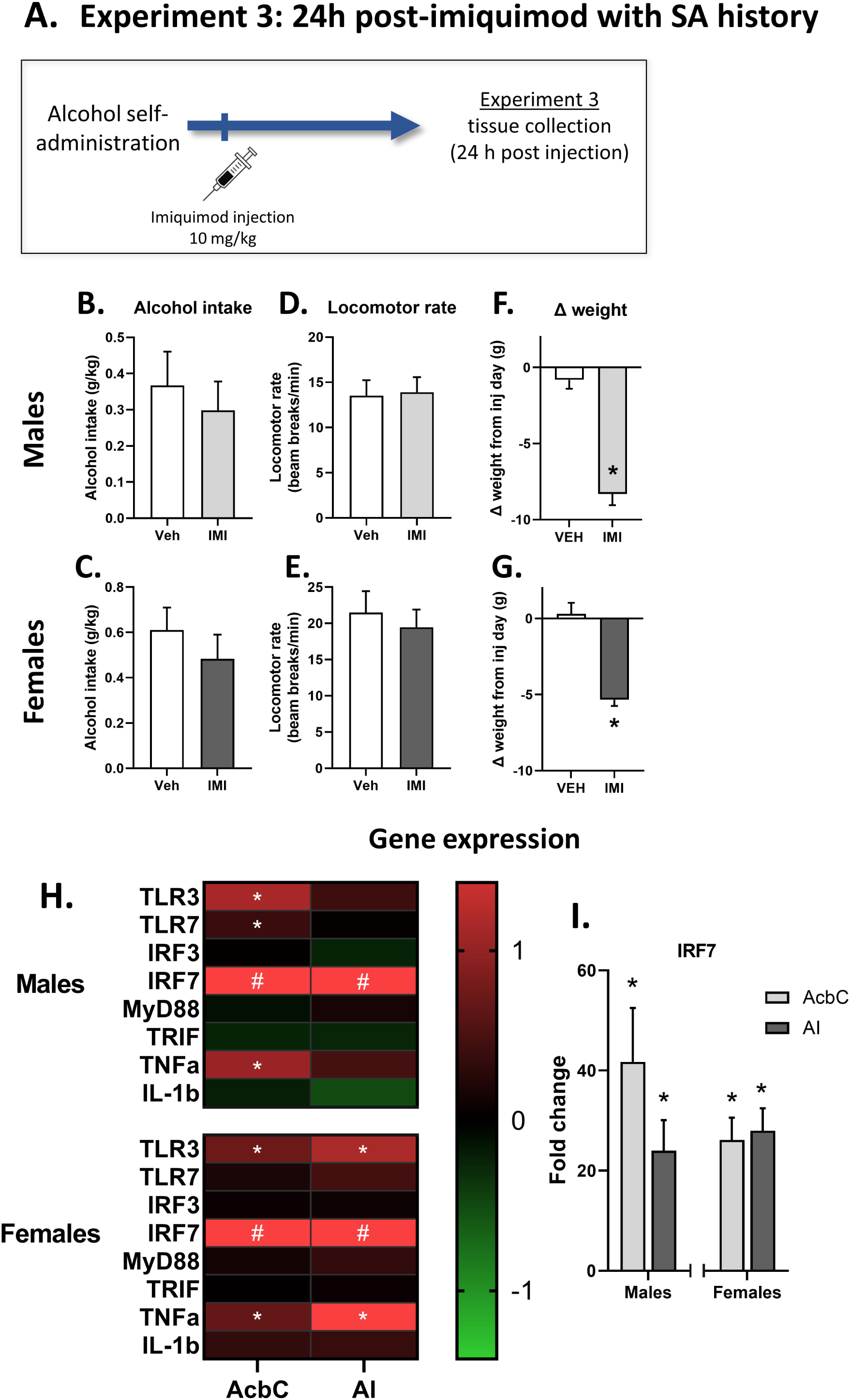
Experiment 3 behavior and gene expression. **A)** For Experiment 3, rats trained on ethanol self-administration were given an injection of imiquimod or vehicle and tissue was collected 24h post-injection. In experiment 4 rats were given injections before daily self-administration sessions four times across 42 days, then tissue was collected 24h after the last injection. **B-E)** Alcohol intake and locomotor rate, and on the day of injection. **F/G)** weight change between the day of injection and the following day. **H)** Experiment 3 gene expression changes in the (AcbC) and AI induced by imiquimod relative to controls visualized as a heatmap. **I)** IRF7 gene expression was greatly increased in both sexes and targets after a single imiquimod injection in rats with a self-administration history. *p<0.05 compared to vehicle control, # p<0.05 compared to control and value is greater than the heatmap scale.

#### Experiment 4: Effects of repeated imiquimod injections on EtOH self-administration and gene expression 24 h post-imiquimod injection

Male and female rats expressing stable self-administration of 15% ethanol after daily self-administration for 24 days were injected with imiquimod (0 or 10 mg/kg, IP; n=12/dose/sex) once every 15 days for a total of 4 injections with 14 self-administration days between each injection (figure 3A). On injection days, imiquimod was administered 2 hours prior to a standard 30-minute self-administration session. The 15 days in-between injections were standard self-administration sessions, as described above. For the figures and analyses the first 3 days post injection are used for analysis as self-administration returned to control levels. The 15 days between injections was used to ensure that there was no residual drug effect. Alcohol intake during the operant session is shown as the primary outcome measurement, and number of active lever presses are shared in supplemental figure 1. Rats were sacrificed 24 hours after the 4^th^ imiquimod injection, spleens were weighed, and brain was collected for RT-PCR analysis.

**Figure 3:**
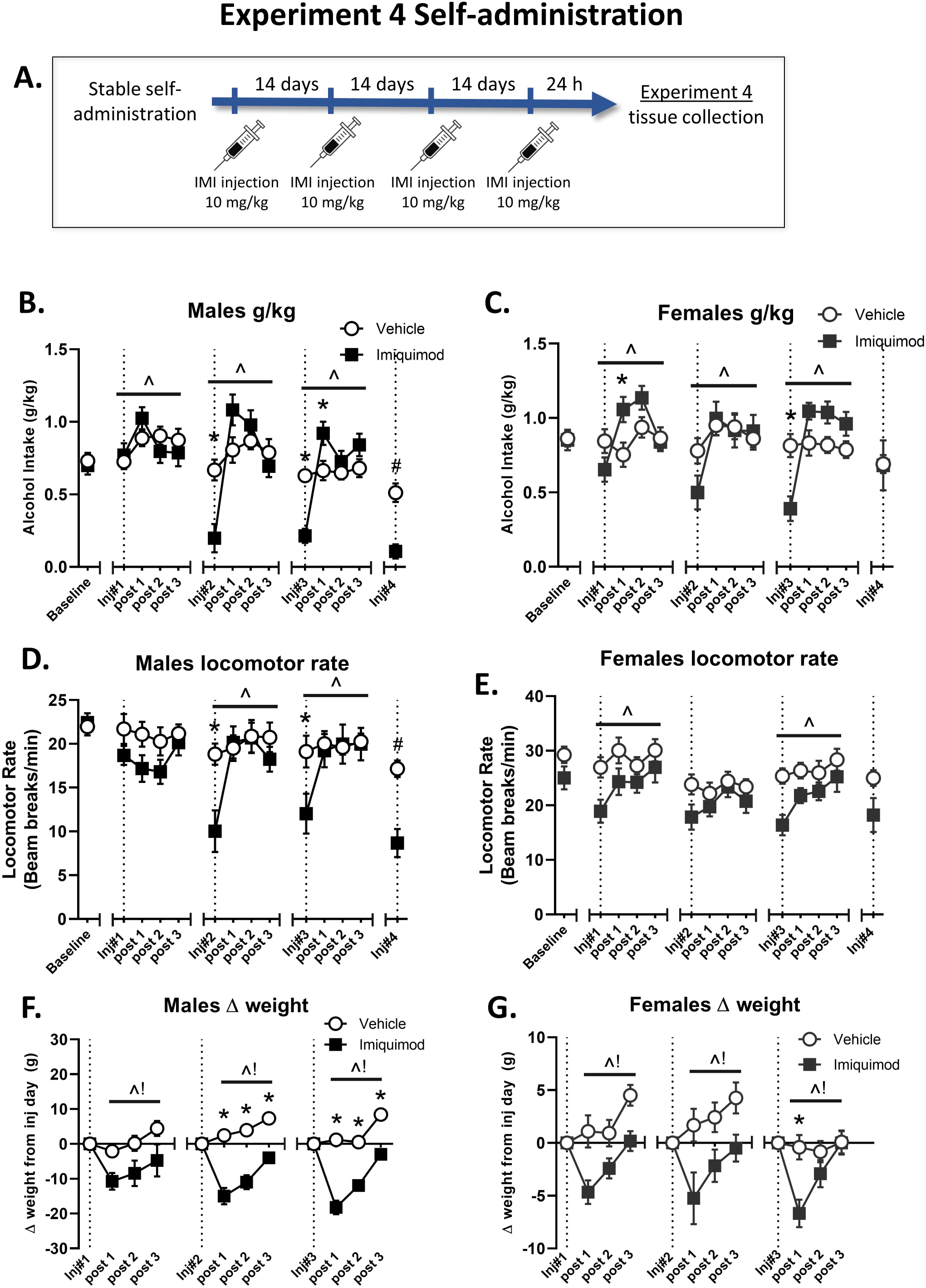
Experiment 4 self-administration behavior. Self-administration at two hours post-imiquimod injection results from Experiment 4. Baseline data are the average of the three training days prior to the first imiquimod injection, and each injection day is represented as a dotted line with subsequent days designated (e.g. post 1 is the following day). **A)** In males, alcohol intake was unaffected by the first injection but subsequent injections reduced alcohol intake on the day of injection and increased intake the day after the third injection. **B)** However in females the first injection resulted in an increase in intake that was not present after subsequent injections, and a reduction in drinking on injection day occurred after the third injection and was not present after the fourth. Like the reductions in drinking, locomotor reductions were present on injection day starting with the second injection in males **(C)** while females did not display similar imiquimod-induced reductions. Finally, imiquimod resulted in weight loss after every injection in both sexes **(E/F)**. *significant interaction and post-hoc difference on day of testing, ^ main effect of test day, ! main effect of imiquimod.

### Statistical analyses

#### Gene expression

We used the 2^ΔΔCt method to determine fold change relative to controls [28]. Fold changes were normalized so that average control fold change equaled 0. For all gene expression and spleen weight data, t-tests were used to determine the effect of imiquimod relative to controls within each sex. Graphs are displayed as expression relative to control subjects with 1 representing the average fold change of controls. Samples were removed from analysis in case of experimenter error or if determined to be a statistical outlier (greater than 2 std. dev. from the mean). All gene expression data including control values are reported as % change from control in Supplemental tables 1-4.

#### Behavioral analyses

In order to take into account IMI induced weight changes, our primary measure in the self-administration studies was alcohol intake, calculated from rat body weight and the number of reinforcers delivered. Alcohol lever responses are presented in Supplemental Figure 1. For self-administration, alcohol lever responses, estimated alcohol intake (g/kg; calculated based on rat body weight and number of reinforcers delivered) and locomotor rate were analyzed using one-way repeated measures analysis of variance (RM-ANOVA) for each sex. Post-hoc analysis (Tukey) was used to determine differences between specific days of training. For the test days and subsequent three self-administration sessions, all behaviors of interest were analyzed by two-way repeated measures ANOVAs for each injection day and the proceeding three days in order to determine the immediate and short-term effects of imiquimod on behavior. Behavior on the fourth and final injection day was analyzed via t-test.

## Results

### Experiment 1: Central gene expression 2 h post-imiquimod injection (figure 1 B and C)

This study investigated the response to the small molecular selective TLR7 agonist imiquimod, in both males and females 2 h after injection. The gene expression (fold change) analysis is illustrated by the heatmap in Figure 1b (see supplemental table 1 for data values). *<u>AcbC</u>*. At 2h-post IMI, IRF7 was greatly increased in both males (9.4-fold; t(11)=7.38, p<0.01) and females (5.3-fold; t(14)=3.13, p<0.01; figure 1C). TLR3 gene expression was elevated 1.7-fold following IMI only in females (t(11)=4.04, p<0.01). *<u>AI</u>*. In contrast, in the AI TLR3 expression was increased 1.7-fold only in males (t(12)=2.77, p<0.05). IRF7 was again greatly increased in both males (2.8-fold; t(12)=2.99, p<0.05) and females (5.3-fold; t(11)=6.88, p<0.0001; figure 1C). MyD88 was increased in females only in the AI (1.5-fold; t(13)=2.77, p<0.05).

### Experiment 2

The gene expression (fold change) analysis is illustrated by the heatmap in Figure 1e (see supplemental table 2 for data values). *Central gene expression 24 h post-imiquimod injection. <u>AcbC</u>*. 24h post-imiquimod, TLR3 gene expression was increased the AcbC in both males (2.2-fold: t(14)=3.03, p<0.01) and females (1.9-fold: t(12)=4.83, p<0.001), while TLR7 expression was increased specifically in males (1.4-fold: t(12)=2.41, p<0.05; see figure 1E). Similar to Experiment 1, IRF7 expression was greatly increased in both brain regions in males (11.6-fold: t(10)=13.69, p<0.0001) and females (12.2-fold: t(12)=4.58, p<0.01; figure 1F). *<u>AI</u>*. In the AI both males and females had increased expression of TLR3 (male 1.8-fold: t(14)=2.70 p<0.05; female 1.6-fold: t(13)=2.32, p<0.05), while like in the AcbC TLR7 expression was increased only in males (1.8-fold: t(14)=2.27, p<0.05; figure 1E). IRF7 gene expression was once again greatly increased in the AI in both males and females (males 5.5-fold: t(10)=3.13, p<0.05); females 3.8-fold: t(10)=2.26, p<0.05; figure 1F). These findings suggest that a single dose of IMI induces IRF7 mRNA, a direct downstream transcription factor target of TLR7, in both AcbC and AI at 2 hrs. that persists for at least 24hr, whereas IRF3, TNFα or IL-1β are not altered. These findings are consistent with the small molecule IMI activating brain TLR7 receptors.

**Table 2.**
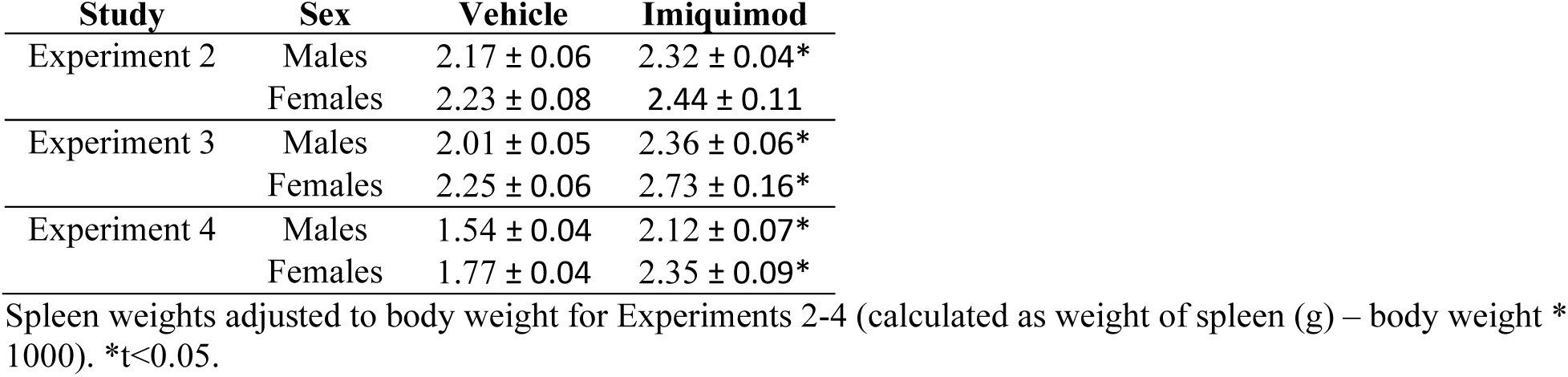
Spleen weights.

Increased spleen weights are commonly used to assess systemic immune responses. At 24 hr. imiquimod increased spleen weight relative to body weight in males as compared to controls (table 2; t(14)=2.18, p<0.05), indicative of induction of an inflammatory response. Female spleen weights were more variable, showing a similar magnitude in spleen weight increase that was not statistically significant (p=0.158). Overall these findings suggest that imiquimod induced a strong inflammatory response.

### Experiment 3: Behavior and weights

During the self-administration session on the imiquimod injection day, neither male nor female rats reduced their drinking nor were locomotor rates affected (figure 2 B-G). However, the following day both males (t(18)=7.87, p<0.0001) and females t(18)=6.52, p<0.0001) had lost weight indicating that imiquimod induced sickness (figure 2 F,G).

### Effect of EtOH self-administration history on gene expression 24 h after a single imiquimod injection (figure 3J and K). AcbC

In rats with a history of alcohol self-administration TLR3 gene expression was increased in the AcbC in both males (2.4-fold; t(18)=3.49, p<0.01) and females (1.7-fold; t(17)=3.42, p<0.01). A small but significant 1.4-fold increase in TLR7 was found only in males (t(17)=2.23, p<0.05). Imiquimod induced expression of the pro-inflammatory cytokine TNFα in males (1.9-fold; t(16)=2.84, p<0.05) and females (1.6-fold; t(17)=2.88, p<0.01). IRF7 gene expression was greatly increased in both males (42.7-fold; t(18)=3.88, p<0.01; figure 2D) and females (27.2-fold; t(17)=5.57, p<0.0001). *<u>AI:</u>* In the AI TLR3 gene expression was increased 2-fold specifically in females (t(18)=3.30, p<0.01). IRF7 expression was again greatly increased in both sexes (males 25-fold; t(18)=4.19, p<0.001 females 29-fold; t(19)=5.92, p<0.0001). Again, expression of TNFα was increased specifically in females (2.3-fold; t(16)=3.61, p<0.01). See supplemental table 3 for data values.

### Spleen weights

Imiquimod resulted in increased spleen weights in both males (t(18)=3.97, p<0.001) and females (t(17)=2.90, p<0.01; table 2).

### Experiment 4: Effects of repeated imiquimod injections on EtOH self-administration 24 h post-imiquimod

#### Self-administration (figure 3):1^st^ injection

During the first injection cycle, two-way repeated measures ANOVAs found main effects of test day in males in alcohol intake [F(3, 66)=3.87, p<0.05; figure 3B] and change in weight [F(2, 44)=18.80, p<0.05; figure 3D], and imiquimod resulted in weight loss [F(1,22)=4.771, p<0.05; figure 3F]. In females there was a significant interaction in alcohol intake [F(3,66) = 5.30, p<0.01; figure 3C] and post-hoc analysis found that imiquimod induced increased drinking the day after the injection (p<0.05). There were significant main effects of test day in both locomotor rate [F(3,66)=6.12, p<0.001; figure 3E] and change in weight [F(2,44)=12.12, p<0.001], and as was the case in males imiquimod resulted in weight loss in females [F(1,22)=12.16, p<0.01; figure 3G].

#### 2^nd^ injection

After the second injection, in males there were significant interactions in both alcohol intake [F(3,66)=7.88, p<0.001; figure 3B] and locomotor rate [F(3,66)=4.73, p<0.01; figure 3d] with imiquimod decreasing both on the day of the injection (p’s<0.01 and 0.05, respectively). There was also a significant interaction in change in weight [F(2,44)=6.86, p<0.01; figure 3F] with imiquimod decreasing weight on all three days post injection (p’s<0.001). Females did not show behavioral decreases on the day of injection as only main effects of test day were seen in alcohol intake [F(3,66)=5.68, p<0.05; figure 3C] and change in weight [F (2,44)=7.62, p<0.01; figure 3G], and imiquimod generally resulted in weight loss [F(1,22)=6.73, p<0.05].

#### 3^rd^ injection

After the third injection, there was a significant interaction of imiquimod and day of testing in males [F(3,66)=17.32, p<0.001; figure 3B] with rats drinking less on injection day (p<0.001) and more on the first day post-injection (p<0.05). There was also a significant interaction in locomotor rate [F(3,66)=3.24, p<0.05; figure 3D] with imiquimod reducing general activity on the day of injection (p<0.05). Lastly, there was an interaction in weight change as well with imiquimod resulting in reduced weight on all post-injection days (p’s <0.001). In females there was also an interaction in alcohol intake [F(3,66)=11.45, p<0.001; figure 3C] with females drinking less on the day of injection (p<0.001). There was also a main effect of test day in locomotor rate [F(3,66)=7.95, p<0.001; figure 3E]. Lastly, a significant interaction in weight change [F(2,44)=6.72, p<0.01; figure 3F revealed significant weight loss on the day after the injection only (p<0.001).

#### 4^th^ injection

On the last injection day, t-tests found decreases in alcohol intake (p<0.001) and locomotor rate (p<0.001) in males only, with imiquimod having no effect in females (figure 3).

### Effects of repeated imiquimod injections on gene expression 24 h post-imiquimod (figure 4B)

#### AcbC

In rats trained to self-administer alcohol, 24h after four repeated imiquimod injections TLR3 was increased in both males (2.6-fold; t(19)=3.15, p<0.01) and females (2.1-fold; t(21)=2.89, p<0.05). TLR7 gene expression was increased 1.9-fold specifically in females (t(22)=4.73, p<0.001) but the increase did not reach the level of significance in males (p=0.057). As has been seen in all of our experiments, IRF7 gene expression was greatly increased in both males (23.4-fold; t(17)=6.56, p<0.0001) and females (16.5-fold; t(19)=4.73, p<0.0001; figure 4D). MyD88 was also increased specifically in males in the AcbC (t(19)=2.32, p<0.05). *<u>AI</u>*. TLR3 gene expression was increased in the AI in both males (1.6-fold; t(19)=3.02, p<0.01) and females (2.1-fold; t(19)=2.77, p<0.05). As was seen in the AcbC, TLR7 gene expression was increased specifically in females (1.5-fold; t(20)=2.39, p<0.05)) but not in males. Lastly, IRF7 was greatly increased in both sexes (males 15.5-fold; t(19)=6.31, p<0.0001: females 18.7-fold; t(21)=6.20, p<0.0001; figure 4C). See supplemental table 4 for data values.

**Figure 4:**
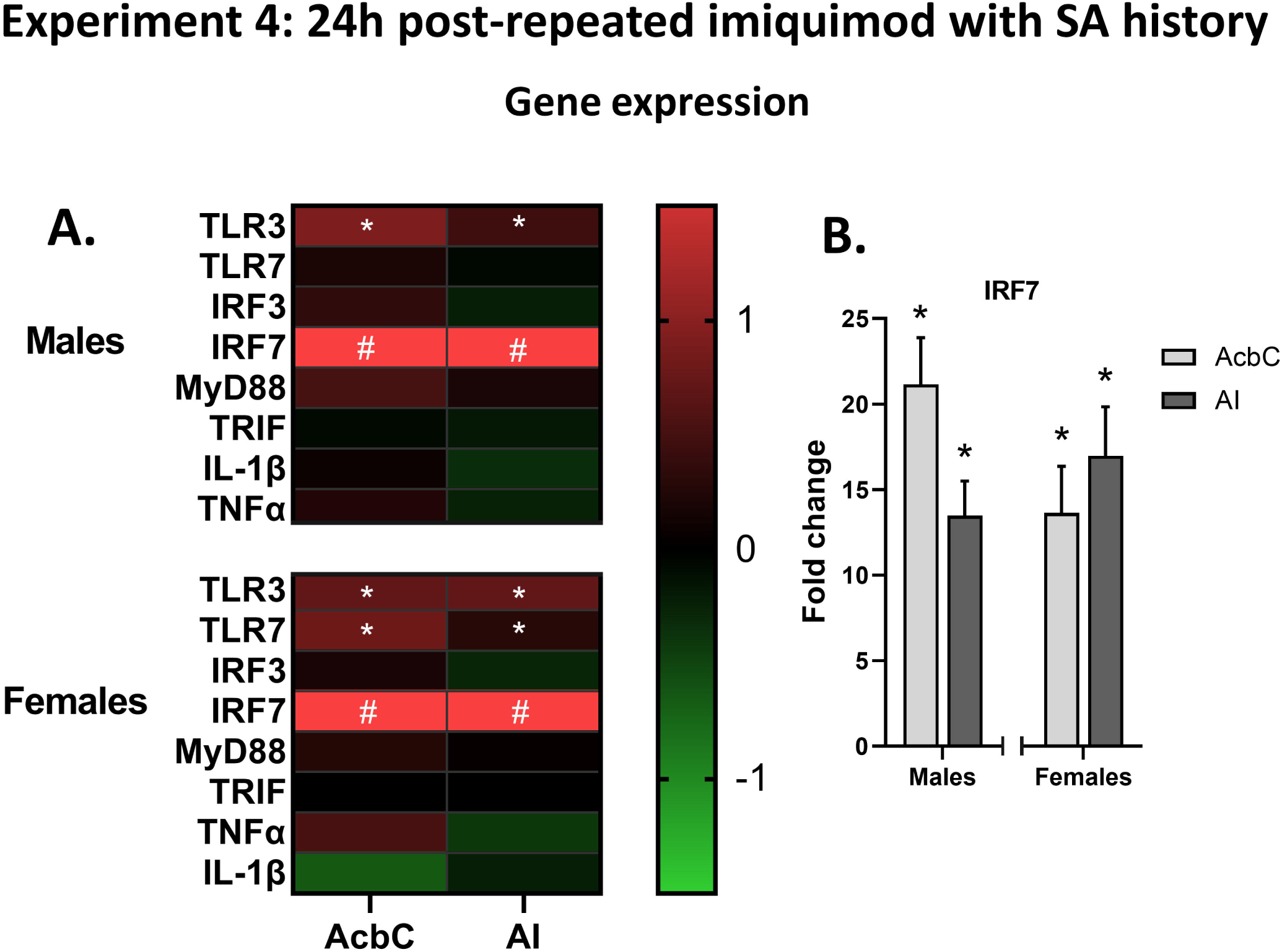
Experiment 4 gene expression. **A)** Experiment 4 gene expression changes in the (AcbC) and AI induced by imiquimod relative to controls visualized as a heatmap, showing consistent increases in TLR3 and IRF7 across brain regions and sex. **B)** Multiple injections of imiquimod resulted in greatly increased expression of IR7 in both brain regions and sexes. *p<0.05 compared to vehicle control, # p<0.05 compared to control and value is greater than the heatmap scale.

### Spleen weights

Imiquimod resulted in increased spleen weights in both males (t(7.51), p<0.0001) and females (t(5.78, p<0.0001; table 2).

## Discussion

The role of TLR7 signaling in operant alcohol self-administration behavior has not been well-established, thus we examined gene expression changes induced by administration of the TLR7 receptor agonist imiquimod and its effects on operant self-administration in male and female rats. In naïve rats, we first determined what genes are modulated by TLR7 activation in the AcbC and AI two brain regions previously implicated in alcohol seeking and self-administration [13, 29-32]. In both sexes, we observed a profound induction of IRF7 in both brain regions at 2 and 24h as well as increases in expression of TLR3 and 7, particularly at 24h. In male and female rats with a self-administration history, a single injection of imiquimod did not significantly reduce alcohol drinking 2 h after injection (figure 2). 24 h later, this group showed an increase in IRF7 that was higher than in the alcohol naïve rats and again increases in TLR3 and 7 genes were observed, as well as an increase in the pro-inflammatory cytokine TNFα. Finally, in the last experiment we examined the consequences of multiple imiquimod injections on alcohol self-administration, and found that males were largely insensitive to the first injection but imiquimod increased drinking the day after subsequent injections, whereas females showed increased intake on the day following the first injection which was reduced after subsequent injections (figure 3). Interestingly, after four imiquimod injections IRF7 and TLRs 3 and 7 expression were increased, but TNFα was not, suggesting the possibility of a molecular adaptation.

After acute administration of imiquimod in naïve rats, at 2 h post-injection we found that TLR3 gene expression was increased in the AI in males, but in females the increase was in the AcbC (figure 1B). By 24 h TLR3 was increased in both brain regions in both sexes, and TLR7 expression was significantly elevated in the males as well (figure 1C). The sex and region-specificity of TLR3 activation at the 2 h timepoint followed by consistent activation across all groups at 24h may be suggestive of sex- and brain region-specific time course of immune induction. With the TLR7 agonist in the present work we found a similar pattern in the AI in females and in the AcbC in males. Further studies will be needed to determine how the neuroimmune response differs across the factors of immune-inducing agent, brain region, and sex. TLR signaling, and specifically TLR3 and TLR7 activation, has been shown to induce microglial activation and chemotaxis [4, 33] and can modulate neuronal morphology, activity and synaptic plasticity [14, 34, 35]. TLR7 is primarily expressed in microglia whereas TLR3 is expressed in neurons, astrocytes, and microglia [36-38], thus the induction of TLR7 only in males at 24 h may be indicative of microglial activation or priming specifically in the males that is not present in females. Alternatively, the time course of TLR7 activation may not be adequately captured by the study’s timepoints, either increasing and resolving between 2 and 24 h or delayed past 24 h. IRF7 expression was increased by imiquimod across all of the 2 and 24 h timepoints and in both brain regions, in agreement with findings after administration of the TLR7 agonist R848 in male mice [12]. IRF7 signaling is thought to be downstream of MyD88 [14], thus it is surprising that MyD88 was increased only in the AI in females at 2h post-injection. However, in an experiment where TLR7 agonist R848 was administered, MyD88 was increased in male mice at 8h post-injection but was resolved to baseline at 24h (in agreement with our findings), and female mice have yet to be examined. In fact, in that work, nearly all TLR-related targets were elevated at 8h (TLR7, TLR4, MyD88, IRF7, TRIF, IRF3) and resolved by 24h save for IRF7, suggesting that the 2h time point used in the present work may be too early for detection of peak immune induction of many of our targets which have previously been shown to be important in affecting alcohol intake in mice [10, 11]. This time point was chosen as imiquimod has a half-life of 2 hours [39] and has been shown to induce fever by 2h at a lower dose of 5 mg/kg [40], but it may be that some aspects of inflammatory gene expression peak significantly later. Indeed, with said lower dose of imiquimod, TNFα and IL-6 expression were increased at 6 h but not 2 h in the rat hypothalamus [40], so the peak immune response may occur later than the peak blood concentration of imiquimod. Additionally, it is possible that rats and mice differ in the time course of the response to TLR7 activation, and the TLR7 agonists used did differ between our study and the Warden et al. studies [10, 11], however the finding that IRF7 remained elevated 24 h post-injection in both species is an interesting consistent finding. Lastly, in the 24 h group we did not find changes in pro-inflammatory cytokines TNFα or IL-1β in either brain region. We did not examine pro-inflammatory cytokines at the 2 h timepoint in experiment 1 as that timepoint was chosen to target TLR signaling molecules rather than cytokines.

In rats that had an alcohol self-administration history we found that acute imiquimod 2 h before the session did not significantly alter alcohol intake nor locomotor rate (Experiment 3, figure 2B-E). However, imiquimod did likely induce sickness as rats lost weight overnight and had enlarged spleens at time of sacrifice 24 h later (figure 2 F/G; table 2), both of which are indicative of induction of an immune response. The timepoint post-injection is the same between Experiments 2 and 3, and though we do not directly compare the two studies, it appears that the alcohol history resulted in a greater IRF7 induction (figure 1E vs. figure 2I; 20-40-fold vs. 13-20-fold, respectively). As has been well-established in the literature, microglia can be primed by an initial stress or pharmacological challenge, including alcohol, resulting in a stronger response to a subsequent challenge [35, 41-43]. Thus, it may be that the alcohol self-administration history primed microglia resulting in enhanced neuroimmune response to imiquimod. Additionally, IRF7 itself is thought to have a role in priming the immune response, so it may be that the history of alcohol self-administration induced changes in microglia that led to a sensitized IRF7 response after imiquimod challenge [44]. There is some precedent for alcohol history to result in increased expression of IRF7 after chronic alcohol administration in adolescent rats [9], further study will need to be done to determine whether a similar response occurs in adults and the alcohol dose range and administration schedules that would promote such priming effects.

Curiously, while the IRF7 response appeared to be heightened in both brain regions in both sexes consistent with what was found in naïve rats, the TLR expression was not as consistently elevated 24h after imiquimod in these rats with a self-administration history. In males TLRs 3 and 7 were elevated in the AcbC but were not increased in the AI, while in females TLR3, but not TLR7, was increased in both brain regions (Figure 2H). These adaptions may be due to induction of molecular tolerance due to the history of alcohol self-administration. Previous studies have found reduced expression of these targets with repeated administration of immune challenge as compared to a single dose [12], and alcohol may be functioning similarly in a sex-and region-specific manner. As has been seen previously imiquimod and R848 TLR7 activation *in vitro*, central gene expression of pro-inflammatory cytokine TNFα was increased in both brain regions in females and in the AcbC in males [36, 45]. Notably, we did not see increased TNFα in alcohol naïve rats in Experiment 2, suggesting that the alcohol history primed the neuroimmune response resulting in greater pro-inflammatory cytokine expression.

As an effect on drinking was not observed 2 h following imiquimod injection in Experiment 3, Experiment 4 was designed to more thoroughly characterize imiquimod effects on self-administration behavior (figure 3A). For simplicity, first we will discuss self-administration on the day of injection (depicted by dotted lines). Consistent with Experiment 3, there were no changes in drinking 2 hours following imiquimod in males or females (figure 3 B,C). However, following the second, third and fourth injection, males greatly reduced their drinking 2 h post-imiquimod. In females, reduced drinking was observed 2 h after the third injection. Interestingly, on the fourth and final injection day drank the same amount of alcohol as controls which suggests sensitization followed by adaptations that result in tolerance to TLR7 agonism. Notably this adaptation was not seen in males. In males, reduction in locomotor activity 2 h following the imiquimod injection mirrored the reductions in drinking, suggesting that both effects may be due to the induction of sickness behavior (figure 4D). In females this comparison did not reach the level of significance though the pattern visually appeared similar (figure 4E). Notably, every injection resulted in substantial weight loss in both sexes (figure 3 F,G) which may suggest that tolerance in respect to behavior, molecular expression, and sickness are not correlated, are not necessarily occurring in the same time frame, or in some cases may not develop at all.

The day following the first imiquimod injection females drank significantly more alcohol as compared to controls while males did not. Subsequently, females did not display that increase with repeated injections while males did, drinking more than controls after the third injection. This is somewhat consistent with the other existing experiment that found increased home cage alcohol intake after administering a TLR7 agonist (R848) for 10 days every other day. After a two-week incubation period mice were tested using an every other day drinking in the dark (EODID) model and exhibited a sustained increase in alcohol intake [12]. Though we did not observe sustained increases in drinking, our design differed in that it utilized operant self-administration and drinking sessions occurred 5 days/week. It is important to note differences between the two studies in both the pattern and amount of drinking, as the EODID procedure used by Grantham et al. resulted in consumption of 5-10 g/kg in control rats across 24h every other day, while in the present study control rats consumed approximately 0.7-1.0 g/kg five consecutive days a week. Our lab previously found increased intake after an 18 day incubation period in operant self-administration when male rats were given 3 mg/kg of the TLR3 agonist poly(I:C) [13], but similarly to the findings in the present experiment the increased intake was not sustained. Additionally, a limitation of the present work is that a single dose of imiquimod was assessed, therefore it is possible that prolonged increase in drinking may be observed using a higher dose. It is noteworthy that few studies examining TLR activation have explicitly examined females, though interestingly peak cytokine levels induced by TLR3 activation by poly(I:C) increased drinking in male mice while reducing intake in females in an EODOD model [10, 11]. Sex-specific neuroimmune responses may provide an explanation for the differences in behavior. The TLR7 gene is encoded on the X chromosome and is known to escape X inactivation [25]. This is thought to explain why women, and individuals with XXY chromosomes (Klinefelter syndrome) are more likely to have increased incidence of autoimmune disorders such as systemic lupus erythematosus [26]. Higher initial TLR7 expression levels may explain why in Experiment 4 females were initially more behaviorally sensitive to TLR7 activation than males, showing increases in alcohol consumption after the first injection. However, the gene expression differences between the sexes after 1 and 4 injections do not necessarily match the changes in drinking, making interpretation difficult. We hypothesized that intake would increase the day following imiquimod injection which did occur, albeit inconsistently across injections. A different administration pattern or dose may be more capable of driving increased alcohol intake in operant-self administration. Regardless, it is clear that TNFα and TLR7 adaptations to repeated TLR7 activation are occurring in both sexes in unique manners.

In experiment 4, tissue was collected from subjects after the fourth imiquimod injection. Spleens in animals that received imiquimod were much larger than vehicle controls likely due to the cumulative amount of imiquimod administered plus possible potentiation resulting from a history of drinking. As was seen in all other experiments, IRF7 expression levels were greatly elevated in both brain regions in males and females (figure 4B). However, IRF7 levels were not as highly expressed as observed in the other group that had a self-administration history after a single imiquimod injection (Figure 3I, 12-22-fold vs. 25-40-fold). Molecular tolerance to repeated TLR7 agonism has been reported [12, 36] and may be responsible for this difference, though a direct comparison within a single study is needed for confirmation. In support of this idea, TNFα was not elevated after 4 imiquimod injections (figure 4A) as was observed in Experiment 3 (figure 2H). The pattern of TLR gene expression was different within each sex as well, with males exhibiting increased TLR3 expression in both the AcbC and AI as opposed to just the AcbC, and no changes to TLR7. Females, instead of showing increased TLR3 in both brain regions after a single imiquimod injection, had increased expression of both TLR3 and TLR7.

As the tissue collection timepoint in Experiment 3 matches the first day of drinking in Experiment 4, it is possible that molecular adaptation that occurs between the two timepoints may be related to the differences in behavior. That is, males did not show increased drinking after the first injection when TLRs were and TNFα increased in the AcbC, but did after multiple injections which also induced greater expression of TLR3 in the AI and no longer led to an increase in TNFα. Similarly in females, at the first drinking timepoint when females had elevated intake it is likely that TLR7 gene expression was down while TNFα expression was up, and after repeated injections when drinking did not differ from controls there was no induction of pro-inflammatory cytokines but TLR7 expression was increased in both brain regions. It is possible that TNFα is part of an immune cascade that drives increased drinking in males but suppresses it in females as was seen in prior studies [10, 11]. As reductions in immune responses after TLR7 agonism have been previously reported [36], these changes are not altogether surprising. Brain region-specific actions of TLRs may be playing an important role in modulating behavior as TLRs are known to have roles outside of antigen recognition [14, 34, 35, 46]. The specifics of these changes will have to be teased apart with studies that manipulate TLRs and their downstream signaling molecules across brain regions and sexes to determine whether they may play a causal role in regulating drinking behavior.

## Conclusion

In summary, we have shown that TLR7 activation reliably induces IRF7 gene expression in both males and females. Further, we demonstrated a role for TLR7 activation in modulation of alcohol self-administration with sex-specific differences in the pattern of drinking across repeated injections. It is noteworthy that some dependent measures, such as pro-inflammatory cytokines and TLR gene expression, drinking, and locomotor activity, changed over time with repeated injections while weight loss and increased IRF7 gene expression were always found.

Interestingly, in a prior study with resiquimod a general trend of reduced inflammatory response with repeated injections was found, which fits well with our TNFα results [36]. The cell-type specificity of the adaptions we observed are not yet known, and it is likely that interactions between multiple cell types are responsible for modulation of behavioral output. As TLR7 is primarily expressed in microglia and is known to modulate neuronal development, function, and morphology [14, 34], it will be important to consider changes that occur within certain cell types rather than simply as a whole in a given brain region. Further, this study examined two brain regions whose interactions are known to modulate alcohol seeking [32, 47], thus an important future direction will be understanding how changes in one region affect the other as well as other relevant brain regions. Imiquimod itself is used as an immune inducer, sold as a topical cream for skin conditions [48] and used as adjunctive treatment with vaccines [49] and with cancer treatments [50-52]. These actions are thought to occur through induction of interferons, particularly interferon-alpha. Thus, it is possible that the molecular cascade involved in interferon induction may be more important in regulation of alcohol-related behaviors than that of classic pro-inflammatory cytokines such as IL-1β and TNFα as were examined here. A greater understanding of the molecular underpinnings of how TLR activation can modulate alcohol drinking will be critical in working towards development of treatment strategies for AUD.

## List of Abbreviations

AI: Anterior insular cortex
AcbC: Nucleus accumbens core
AUD: Alcohol use disorder
MyD88: Myeloid differentiation primary response 88
IRF7: Interferon regulatory factor 7
LPS: lipopolysaccharide
TLR: Toll-like receptor
TNFα: Tumor necrosis factor alpha
TRIF: TIR-domain-containing adapter-inducing interferon-β

## Figure captions

**Supplemental Figure 1: Experiment 3 lever presses**. Active lever responses on injection days and three days after the first three injections. **A)** <u>Males</u>: after the first injection 2 way ANOVA found a main effect of day [F(3,66)=15.43, p<0.0001]. or the second injection there was a main effect of day [F(3,66)=15.42, p<0.0001] and an interaction [F(3,66)=7.08, p<0.001] with imiquimod different from controls on the injection day (p<0.05). Following the third injection there were again significant effects of day [F(3,66)=20.31, p<0.0001] and an interaction [F(3,66)=16.55, p<0.001] with a reduction in lever presses on the injection day (p<0.0001) and an increase the following day (post 1, p<0.01). Imiquimod also reduced lever responses on the final injection day (p<0.0001). **B)** <u>Females</u>: After the first injection there was a significant effect of day [F(3,66)=4.42, p<0.01] and an interaction [F(3,66)=3.95, p<0.05]. After the second injection there was a significant main effect of day [F(3,66)=5.44, p<0.01]. With the third injection there was again an effect of day [F(3,66)=9.68, p<0.001] and an interaction [F(3,66)=9.68, p<0.0001] with post-hoc comparisons finding imiquimod reduced lever responses on the injection day (p<0.01). *significant interaction and post-hoc difference on day of testing, ^ main effect of test day, ! main effect of imiquimod.

## Funding

This work was supported by the National Institute on Alcohol Abuse and Alcoholism (AA025713, AA020024, AA020023, AA011605, AA019767) and the National Institute on Aging (AG072894) of the National Institutes of Health. DFL was supported by F32AA029289 and AA007573.

## Availability of data and materials statement

The datasets used and/or analyzed during the current study are available from the corresponding author on request.

## Authors’ contributions

DFL contributed to design of experiments, collected and analyzed tissue, analyzed and interpreted all results, and was a major contributor in writing the manuscript. WL collected tissue, conducted RTPCR analyses and interpreted results. SEL collected and analyzed behavioral data, collected tissue, analyzed RTPCR results, interpreted results and contributed to writing the manuscript. JL conducted RTPCR analyses and contributed to interpretation of results. KV collected behavioral data, contributed to interpretation of results, and contributed to writing the manuscript. KG contributed to interpretation of results and contributed to writing the manuscript. RPV was a contributor in the design of the experiments and contributed to interpretation of results. FTC was a major contributor in the design of the experiments and contributed to interpretation of results. JB was a major contributor in the design of the experiments, contributed to interpretation of results, and was a major contributor in writing the manuscript.

## Competing interests

The authors declare that they have no competing interests.

## Consent for publication

Not applicable

## Ethics Approval

Experimental procedures were approved by the Institutional Animal Care and Use Committee of the University of North Carolina at Chapel Hill and conducted in accordance with National Institutes of Health regulations for the care and use of animals.

## Notes

### Competing Interest Statement

The authors have declared no competing interest.

