## Supplementary figures and images for "The toll-like receptor 7 agonist imiquimod increases ethanol self-administration and induces expression of toll-like receptor related genes"

### Supplemental Figure 1

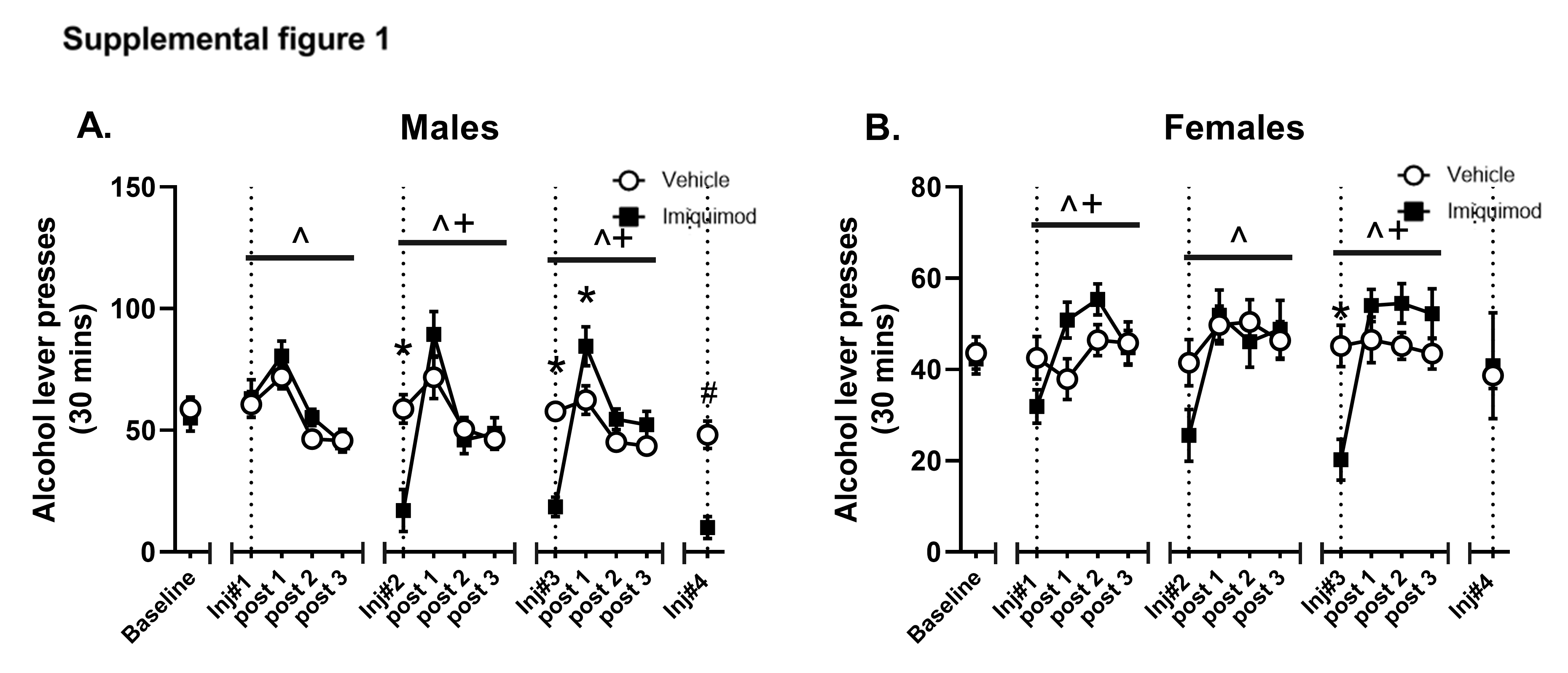
