## Supplemental Tables 1-4 for "The toll-like receptor 7 agonist imiquimod increases ethanol self-administration and induces expression of toll-like receptor related genes"

**Supplemental Table 1**

| Expt 1 | Males | | Females | |
|---|---|---|---|---|
| **AcbC** | **Vehicle** | **Imiquimod** | **Vehicle** | **Imiquimod** |
| TLR3 | 1.00 ± 0.14 | 1.38 ± 0.22 | 1.00 ± 0.09 | 1.70 ± 0.14* |
| TLR7 | 1.00 ± 0.086 | 0.88 ± 0.12 | 1.00 ± 0.15 | 1.10 ± 0.13 |
| IRF3 | 1.00 ± 0.068 | 1.12 ± 0.09 | 1.00 ± 0.09 | 1.00 ± 0.07 |
| IRF7 | 1.00 ± 0.23 | 9.42 ± 1.21* | 1.00 ± 0.33 | 5.28 ± 1.33* |
| MyD88 | 1.00 ± 0.19 | 0.97 ± 0.21 | 1.00 ± 0.20 | 1.23 ± 0.16 |
| TRIF | 1.00 ± 0.22 | 0.80 ± 0.10 | 1.00 ± 0.12 | 1.00 ± 0.13 |
| **AI** | | | | |
| TLR3 | 1.00 ± 0.18 | 1.72 ± 0.18* | 1.00 ± 0.25 | 0.92 ± 0.10 |
| TLR7 | 1.00 ± 0.11 | 1.16 ± 0.11 | 1.00 ± 0.10 | 0.9 ± 0.05 |
| IRF3 | 1.00 ± 0.23 | 0.87 ± 0.21 | 1.00 ± 0.40 | 1.38 ± 0.58 |
| IRF7 | 1.00 ± 0.17* | 2.78 ± 0.49 | 1.00 ± 0.35 | 5.32 ± 0.44* |
| MyD88 | 1.00 ± 0.09 | 0.83 ± 0.09 | 1.00 ± 0.07 | 1.47 ± 0.15* |
| TRIF | 1.00 ± 0.20 | 0.76 ± 0.07 | 1.00 ± 0.24 | 0.85 ± 0.10 |

Experiment 1 Real-time RTPCR results for each sex, brain region, and target expressed as fold change from control. * indicates significant difference from control.

**Supplemental Table 2**

| Expt 2 | Males | | Females | |
|---|---|---|---|---|
| **AcbC** | **Vehicle** | **Imiquimod** | **Vehicle** | **Imiquimod** |
| TLR3 | 1.00 ± 0.12 | 2.23 ± 0.39* | 1.00 ± 0.10 | 1.85 ± 0.15* |
| TLR7 | 1.00 ± 0.07 | 1.44 ± 0.17 | 1.00 ± 0.10 | 1.01 ± 0.13 |
| IRF3 | 1.00 ± 0.16 | 1.14 ± 0.16 | 1.00 ± 0.12 | 0.87 ± 0.10 |
| IRF7 | 1.00 ± 0.26 | 11.59 ± 0.62* | 1.00 ± 0.23 | 12.15 ± 2.08* |
| MyD88 | 1.00 ± 0.06 | 0.96 ± 0.12 | 1.00 ± 0.07 | 1.27 ± 0.13 |
| TRIF | 1.00 ± 0.17 | 1.03 ± 0.18 | 1.00 ± 0.23 | 0.86 ± 0.08 |
| IL-1β | 1.00 ± 0.32 | 0.80 ± 0.13 | 1.00 ± 0.18 | 0.99 ± 0.13 |
| TNFα | 1.00 ± 0.16 | 1.40 ± 0.16 | 1.00 ± 0.13 | 0.95 ± 0.14 |
| **AI** | | | | |
| TLR3 | 1.00 ± 0.17 | 1.78 ± 0.24* | 1.00 ± 0.15 | 1.62 ± 0.21* |
| TLR7 | 1.00 ± 0.22 | 1.76 ± 0.26* | 1.00 ± 0.18 | 0.83 ± 0.14 |
| IRF3 | 1.00 ± 0.07 | 1.07 ± 0.11 | 1.00 ± 0.14 | 1.35 ± 0.18 |
| IRF7 | 1.00 ± 0.25 | 5.48 ± 0.98* | 1.00 ± 0.23 | 3.80 ± 0.86* |
| MyD88 | 1.00 ± 0.11 | 0.56 ± 0.08 | 1.00 ± 0.21 | 1.01 ± 0.25 |
| TRIF | 1.00 ± 0.06 | 1.12 ± 0.14 | 1.00 ± 0.15 | 0.78 ± 0.09 |
| IL-1β | 1.00 ± 0.15 | 0.87 ± 0.15 | 1.00 ± 0.21 | 1.55 ± 0.12 |
| TNFα | 1.00 ± 0.15 | 1.46 ± 0.16 | 1.00 ± 0.1 | 1.36 ± 0.14 |

Experiment 2 Real-time RTPCR results for each sex, brain region, and target expressed as fold change from control. * indicates significant difference from control.

**Supplemental Table 3**

| Expt 3 | Males | | Females | |
|---|---|---|---|---|
| **AcbC** | **Vehicle** | **Imiquimod** | **Vehicle** | **Imiquimod** |
| TLR3 | 1.00 ± 0.09 | 2.35 ± 0.38* | 1.00 ± 0.10 | 1.73 ± 0.18* |
| TLR7 | 1.00 ± 0.08 | 1.42 ± 0.18* | 1.00 ± 0.11 | 1.36 ± 0.18 |
| IRF3 | 1.00 ± 0.12 | 1.21 ± 0.26 | 1.00 ± 0.09 | 1.09 ± 0.09 |
| IRF7 | 1.00 ± 0.13 | 42.73 ± 10.75* | 1.00 ± 0.19 | 27.15 ± 4.44* |
| MyD88 | 1.00 ± 0.10 | 0.87 ± 0.11 | 1.00 ± 0.12 | 1.13 ± 0.09 |
| TRIF | 1.00 ± 0.12 | 0.77 ± 0.08 | 1.00 ± 0.10 | 1.01 ± 0.10 |
| IL-1β | 1.00 ± 0.11 | 1.41 ± 0.24 | 1.00 ± 0.16 | 0.98 ± 0.12 |
| TNFα | 1.00 ± 0.11 | 1.86 ± 0.28* | 1.00 ± 0.11 | 1.59 ± 0.18* |
| **AI** | | | | |
| TLR3 | 1.00 ± 0.10 | 1.31 ± 0.15 | 1.00 ± 0.11 | 2.01 ± 0.26* |
| TLR7 | 1.00 ± 0.13 | 0.95 ± 0.06 | 1.00 ± 0.10 | 1.43 ± 0.19 |
| IRF3 | 1.00 ± 0.15 | 0.71 ± 0.09 | 1.00 ± 0.10 | 1.31 ± 0.17 |
| IRF7 | 1.00 ± 0.16 | 25.02 ± 6.07* | 1.00 ± 0.12 | 28.99 ± 4.47* |
| MyD88 | 1.00 ± 0.13 | 1.12 ± 0.13 | 1.00 ± 0.10 | 1.31 ± 0.12 |
| TRIF | 1.00 ± 0.14 | 0.87 ± 0.10 | 1.00 ± 0.09 | 1.23 ± 0.20 |
| IL-1β | 1.00 ± 0.15 | 0.64 ± 0.09 | 1.00 ± 0.14 | 0.93 ± 0.19 |
| TNFα | 1.00 ± 0.10 | 1.62 ± 0.37 | 1.00 ± 0.17 | 2.31 ± 0.32* |

Experiment 3 Real-time RTPCR results for each sex, brain region, and target expressed as fold change from control. * indicates significant difference from control.

**Supplemental Table 4**

| Expt 4 | Males | | Females | |
|---|---|---|---|---|
| **AcbC** | **Vehicle** | **Imiquimod** | **Vehicle** | **Imiquimod** |
| TLR3 | 1.00 ± 0.11 | 2.59 ± 0.41* | 1.00 ± 0.10 | 2.13 ± 0.34* |
| TLR7 | 1.00 ± 0.06 | 1.34 ± 0.15 | 1.00 ± 0.08 | 1.85 ± 0.18* |
| IRF3 | 1.00 ± 0.15 | 1.29 ± 0.12 | 1.00 ± 0.11 | 1.31 ± 0.19 |
| IRF7 | 1.00 ± 0.14 | 23.41 ± 3.11* | 1.00 ± 0.17 | 16.51 ± 2.98* |
| MyD88 | 1.00 ± 0.10 | 1.59 ± 0.23* | 1.00 ± 0.11 | 1.39 ± 0.24 |
| TRIF | 1.00 ± 0.16 | 1.07 ± 0.19 | 1.00 ± 0.19 | 0.98 ± 0.17 |
| IL-1β | 1.00 ± 0.16 | 1.43 ± 0.32 | 1.00 ± 0.23 | 0.59 ± 0.17 |
| TNFα | 1.00 ± 0.15 | 1.29 ± 0.22 | 1.00 ± 0.14 | 0.95 ± 0.33 |
| **AI** | **Vehicle** | **Imiquimod** | **Vehicle** | **Imiquimod** |
| TLR3 | 1.00 ± 0.10 | 1.55 ± 0.17* | 1.00 ± 0.07 | 2.05 ± 0.38* |
| TLR7 | 1.00 ± 0.21 | 1.09 ± 0.14 | 1.00 ± 0.06 | 1.45 ± 0.20* |
| IRF3 | 1.00 ± 0.23 | 1.04 ± 0.19 | 1.00 ± 0.19 | 1.19 ± 0.32 |
| IRF7 | 1.00 ± 0.20 | 15.49 ± 2.35* | 1.00 ± 0.12 | 18.69 ± 3.37* |
| MyD88 | 1.00 ± 0.08 | 1.09 ± 0.08 | 1.00 ± 0.10 | 1.05 ± 0.16 |
| TRIF | 1.00 ± 0.17 | 0.82 ± 0.13 | 1.00 ± 0.08 | 0.97 ± 0.14 |
| IL-1β | 1.00 ± 0.15 | 0.73 ± 0.25 | 1.00 ± 0.24 | 1.07 ± 0.25 |
| TNFα | 1.00 ± 0.27 | 1.14 ± 0.35 | 1.00 ± 0.20 | 0.56 ± 0.08 |

Experiment 4 Real-time RTPCR results for each sex, brain region, and target expressed as change from control. * indicates significant difference from control.
